# eEF2 improves dense connective tissue repair and healing outcome by regulating cellular death, autophagy, apoptosis, proliferation and migration

**DOI:** 10.1101/2022.12.10.519857

**Authors:** Junyu Chen, Jin Wang, Xinjie Wu, Nils Simon, Camilla I Svensson, Juan Yuan, David A Hart, Aisha S Ahmed, Paul W Ackermann

## Abstract

Outcomes following human dense connective tissue (DCT) repair are often variable and suboptimal, resulting in compromised function and development of chronic painful degenerative diseases. Moreover, biomarkers and mechanisms that guide good clinical outcomes after DCT injuries are mostly unknown. Here, we characterize the proteomic landscape of DCT repair following human tendon rupture and its association with long-term patient-reported outcome. Moreover, the regulatory mechanisms of relevant biomarkers were assessed partly by gene silencing experiments. A Mass-Spectrometry based proteomic approach quantified a large number (769) of proteins, including 51 differentially expressed proteins among 20 good versus 20 poor outcome patients. A novel biomarker, elongation factor-2 (eEF2) was identified as being strongly prognostic of the 1-year clinical outcome. Further bioinformatic and experimental investigation revealed that eEF2 positively regulated autophagy, cell proliferation and migration, as well as reduced cell death and apoptosis, leading to improved DCT repair and outcomes. Findings of eEF2 as novel prognostic biomarker could pave the way for new targeted treatments to improve healing outcomes after DCT injuries.

## Introduction

Human dense connective tissues (DCTs) are matrix-rich tissues which provide vital support and protection to other tissues and organs in the body. However, the reparative process after an injury, especially after injury to DCTs such as tendon, ligament, meniscus and intervertebral disc, can lead to considerable individual variation in long-term patient outcomes. Such variations often result in compromised function, chronic pain and degenerative musculoskeletal diseases for many patients. Knowledge of the regulatory mechanisms responsible for such variability in regenerative outcomes, as well as identification of validated biomarkers that can be used as predictors of DCT healing are still mostly lacking (Abdul Alim et al., 2018, Baranzini et al., 2021, Chen et al., 2021, Grol et al., 2021).

Prior studies in animals have utilized proteomics to delineate the healing of DCT (Anderson-Baucum et al., 2021, Hussein et al., 2018, Fischer et al., 2022, Tchoukalova et al., 2022). However, there is a very limited number of reports examining how the human proteome changes in this process. One human study has compared the proteomes of keloid scars and healthy skin (Ong et al., 2010). The low number of human proteomic studies of DCT healing presumably relate to problems in obtaining large numbers of clinical specimens and subsequently integrating the findings with patient-reported outcomes. However, some aspects of these limitations can potentially be overcome by using recent technological developments in the field of mass spectrometry (MS)-based proteomics which have considerably improved the sensitivity, completeness and quantitative accuracy of this technique for investigating human diseases (Kim et al., 2014, Wilhelm et al., 2014). Our recent findings using MS to analyze the proteomic landscape of human tendon repair in patients with Achilles tendon rupture (ATR) with high quality and sensitivity (Chen et al., 2022) demonstrate the utility of such approaches.

The Achilles tendon is the largest and thickest dense connective tissue in the human body, as well as one of the most commonly injured DCT (Mazzone and McCue, 2002). Patients with ATR also exhibit considerable variability in patient-reported outcomes after injury (Myhrvold et al., 2022). Previous studies indicated that improved patient-reported outcomes after ATR and other DCT injuries are associated with higher production of collagen type I (Col1) (Maffulli et al., 2002, Abdul Alim et al., 2018, Alim et al., 2016). Col1, produced by fibroblast cells (Li and Wang, 2011), is the most abundant protein in these tissues, bringing structure and strength to the matrix of DCTs. Thus, specific biomarkers with strong association to Col1 synthesis would provide a tool to monitor improvement in DCT repair. However, the regulatory mechanisms regarding Col1 production in fibroblast cells during DCT repair remain far from being fully understood (Liu et al., 2016).

To test the hypothesis that a proteomic approach would lead to the identification of novel prognostic biomarkers of early healing efficacy, we aimed to characterize the proteomic components of human DCT repair by combining MS-based proteomics with validated patient-reported outcomes. The present studies also aimed to assess the best prognostic biomarker utilizing tissue biopsies obtained from ATR patients at the time of repair surgery, and with good and poor 1-year clinical outcomes. Further, the regulatory mechanisms involved for the identified biomarkers on Col1 synthesis were explored in primary fibroblast cells and in fibroblast cell lines by gene silencing to knock down the expression of specific molecules. The findings indicate that eukaryotic elongation factor-2 (eEF2) is such a biomarker for prognosis of good outcomes following injury to a DCT.

## Results

### Patient Cohort

To create a tissue atlas of good and poor connective tissue repair, surgical biopsies from a total of 40 patients with ATR were collected, of which 20 patients were assessed to have had a good clinical outcome and the other 20 patients were evaluated as to having a poor clinical outcome according to the 1-year validated patient-reported outcomes (Svedman et al., 2018). Patient characteristics and 1-year patient-reported and functional outcomes are presented in Table 1.

**Table 1.** Patient characteristics and outcome data

The clinical and functional characteristics of ATR patients included in study.
| Variables |  | Group | Good outcome | Poor outcome | p-value |
| --- | --- | --- | --- | --- | --- |
| N (M:F) | 32:8 | 40 | 20 | 20 |  |
| Age (years) |  | 40 (26,65) | 40 (26,65) | 39 (28,59) | ns |
| BMI (kg/m<sup>2</sup>) |  | 25.4 (19.6,34.7) | 24.6 (19.6,34.7) | 25.4 (20.6,30.6) | ns |
| TTS (hours) |  | 85.8 (24.5,158.7) | 87.5 (24.5,158.7) | 85.3 (41.0,135.6) | ns |
| ATRS |  | 76 (40,100) | 95 (87,100) | 63 (40,78) | <0.001 |
|  | Limitation in calf strength | 8 (1,10) | 9 (7,10) | 5 (1,8) | <0.001 |
|  | Tiredness in the calf | 8 (3,10) | 10 (7,10) | 5 (3,8) | <0.001 |
|  | Stiffness in the calf | 8 (2,10) | 10 (5,10) | 5 (2,8) | <0.001 |
|  | Pain in the calf | 10 (3,10) | 10 (9,10) | 9 (3,10) | <0.001 |
|  | Activity of daily life | 9 (4,10) | 10 (9,10) | 8 (4,10) | <0.001 |
|  | Walking on uneven surface | 10 (3,10) | 10 (7,10) | 8 (3,10) | <0.001 |
|  | Limitation on walking in stairs | 9 (4,10) | 10 (9,10) | 8 (4,10) | <0.001 |
|  | Limitation on running | 8 (0,10) | 10 (8,10) | 6 (0,9) | <0.001 |
|  | Limitation on jumping | 7 (2,10) | 10 (7,10) | 4 (2,9) | <0.001 |
|  | loss in physical work | 9 (3,10) | 10 (9,10) | 8 (3,10) | <0.001 |
| FAOS |  |  |  |  |  |
|  | Pain | 100 (69,100) | 100 (89,100) | 93 (69,100) | 0.007 |
|  | Symptom | 93 (39,100) | 100 (75,100) | 89 (39,100) | 0.009 |
|  | Activity of daily life | 100 (88,100) | 100 (91,100) | 98 (88,100) | 0.012 |
|  | Sport and recreation | 88 (45,100) | 100 (75,100) | 75 (45,90) | <0.001 |
|  | Foot-and ankle-related QOL | 69 (31,100) | 91 (69,100) | 56 (31,88) | <0.001 |
| HRT |  |  |  |  |  |
| (LSI%) | Concentric power | 80.8 (23.5,189.9) | 80.6 (52.9,115.9) | 82.1 (23.5,189.9) | ns |
|  | Total work | 71.5 (24.3,288.0) | 77.7 (57.5,119.6) | 68.2 (24.3,288.0) | ns |
|  | Repetition | 90.5 (45.8,275) | 96.7 (71.4,114.3) | 82.7 (45.8,275.0) | ns |
|  | Average height | 81.8 (37.6,110.5) | 81.8 (65.1,110.5) | 81.9 (37.6,104.0) | ns |

### Quantitative Proteomic Characterization of Good and Poor Outcome Patients

To identify proteins that are potentially prognostic of good and poor Achilles tendon repair, proteins from the tissue samples were extracted and subsequently separated using reverse phase liquid chromatography. The proteome of the extracted proteins was developed using mass spectroscopy as detailed in the methods section. The proteomic data was then grouped based on the 1-year clinical outcomes for the patients (Svedman et al., 2018). The computational data analysis detected 855 unique proteins, including 769 shared proteins across the good and poor outcome groups (Figure 1a). Among the shared proteomic file, 51 differentially expressed proteins were identified with 10 down- and 41 up-regulated proteins in good when compared with poor outcome subgroup (Figure 1b). By analysis of enrichment factor, it was observed that the most enriched processes for the down-regulated proteins included myofibril assembly and skeletal tissue development. The up-regulated proteins were mostly involved in extracellular matrix (ECM) organization, collagen metabolism, and inflammatory and immune responses (Figure 1c).

**Figure 1.**
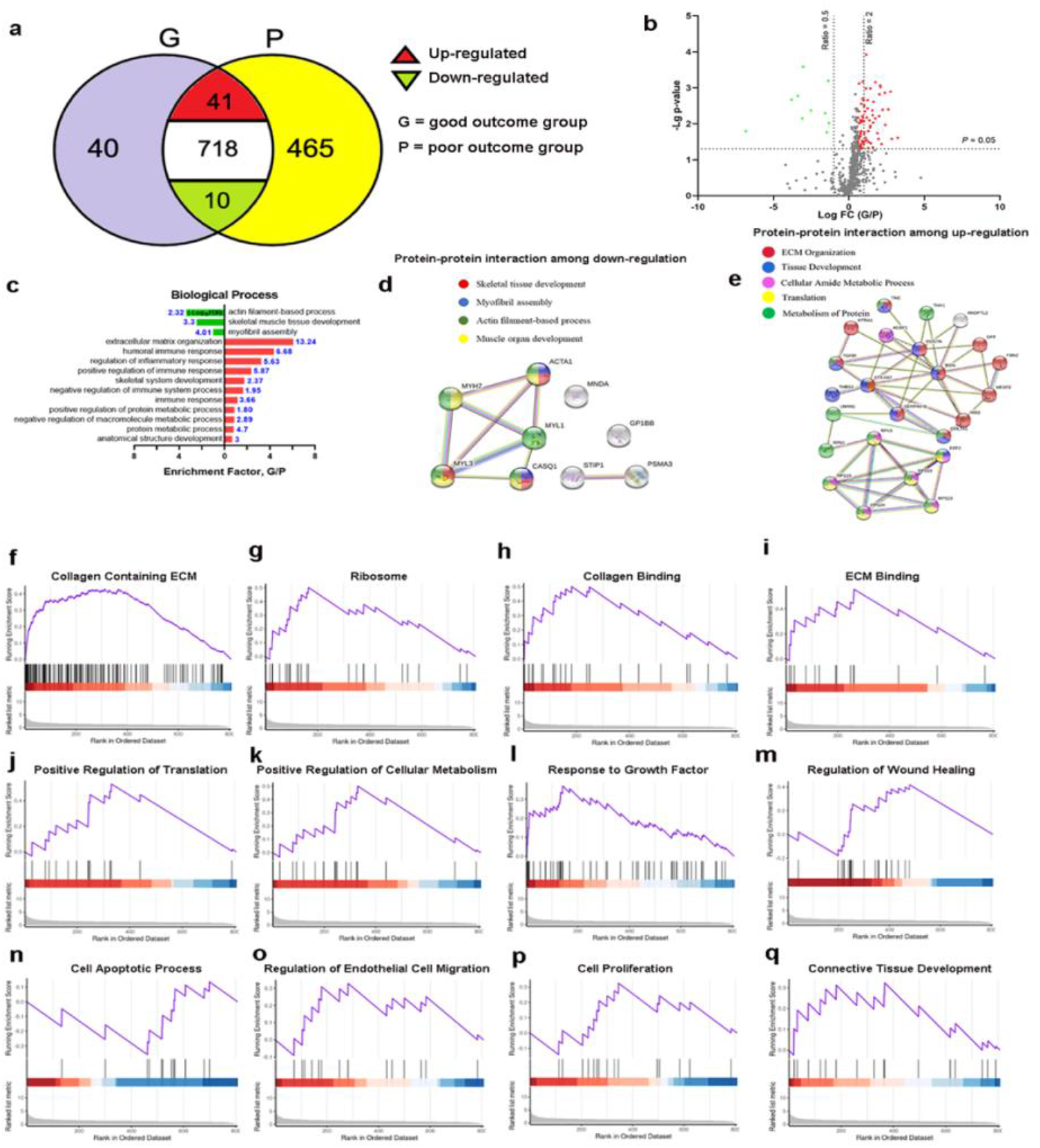
Quantitative proteomic file of injured human Achilles tendon. a) Venn plot of overlapping and distinct proteomes of patients with good (G) and poor (P) outcome. n = 20 in each group; b) Volcano diagram with differentially expressed proteins. The X coordinate represents Log_2_ fold change (FC) and the Y coordinate to −Log_10_ (*p*-value). Each dot represents a protein with red = up-regulated, green =down-regulated and, black = non-differentially expressed proteins; c) Bar plot of differentially expressed proteins with matched biological processes. Size of bar represents the number; −log10 FDR was used as reliability, red = up-regulated while green = down-regulated proteins; Enrichment factor shows the ratio of pathway related proteins to the whole identified proteome. Protein-protein interactions among d) down-regulated, and e) up-regulated proteins. STRING v11.0 was used to analyze functional enrichment. Relevant biological processes are presented with different colors; f-q) GSEA plots of the highly involved pathways related to differentially expressed proteins.

A highly enriched protein-protein interaction among the 10 down-regulated proteins was detected, highlighting potential biological processes of skeletal tissue development, myofibril assembly and muscular development (Figure 1d). The protein-protein interaction among 41 up-regulated markers was also identified, which revealed that these proteins are mainly involved in ECM organization, tissue development and protein metabolism (Figure 1e). GSEA of the 51 (10 down- and 41 up-regulated proteins) differentially expressed proteins detected collagen binding, ECM regulation, metabolic pathways, as well processes involved in wound healing, cell migration and proliferation, and connective tissue development as potential enriched pathways (Figure 1f-q).

### Detection of Predictive Biomarkers for Dense Connective Tissue Repair

To identify potential prognostic biomarkers, a multiple-regression analysis using the proteomic and clinical and functional data was performed. As a result, two up-regulated proteins, eEF2 (Figure 2a-c) and fibrillin-2 (FBN2) (Figure 2d-f) were identified, both of which were positively associated with improved ATRS and average heel rise height. However, only eEF2 with an area under the curve (AUC) value of 0.86 (> 0.85), as compared to FBN2 with an AUC of 0.69, exhibited a strong prediction of good clinical outcome (Figure 2g).

**Figure 2.**
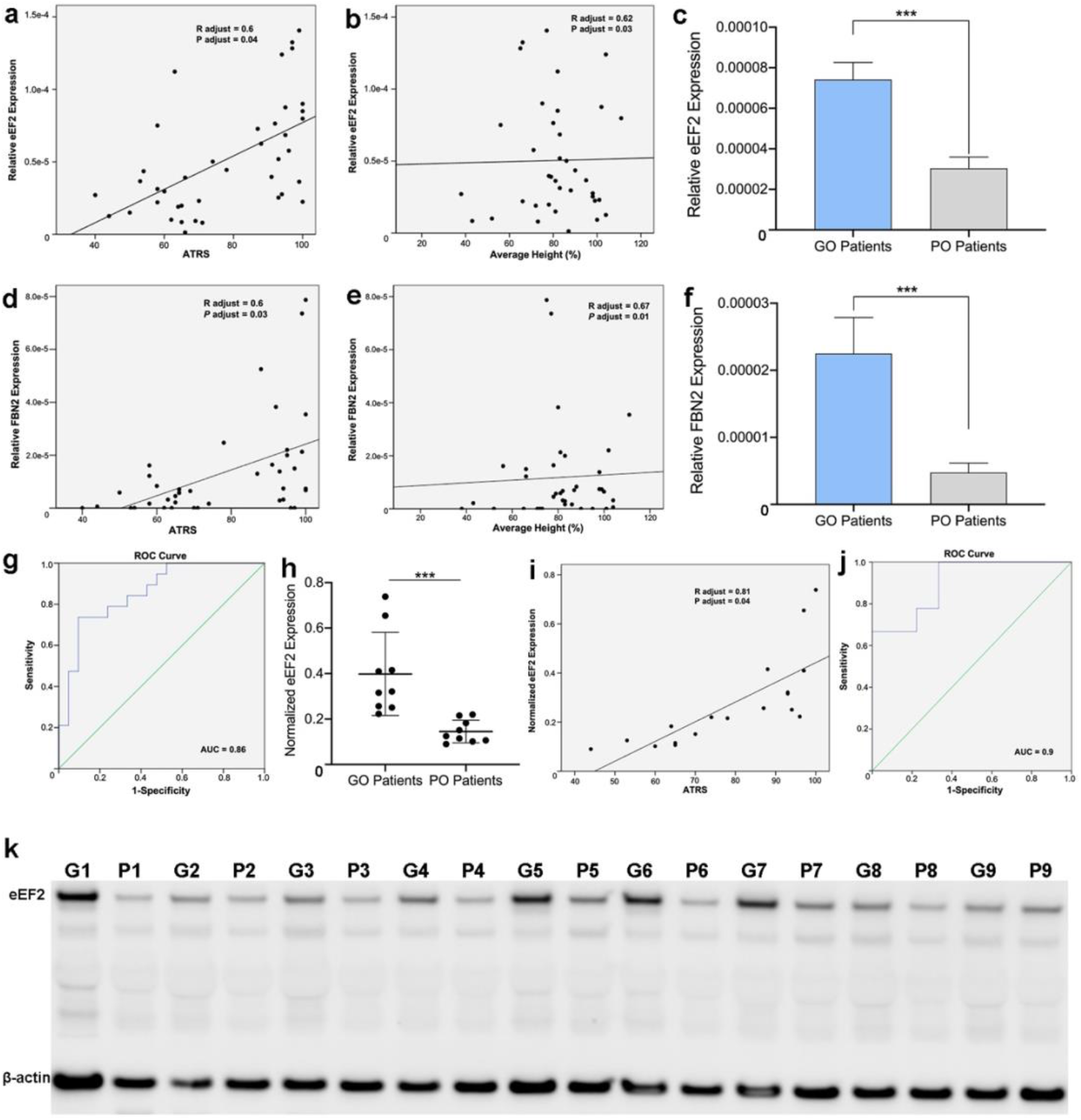
Prognostic biomarker selection and verification in injured Achilles tendon tissues. Association of eEF2 expression with a) ATRS and, b) average heal rise, n = 40; c) eEF2 expression between good outcome (GO, n = 20) and poor outcome (PO, n = 20) patients, data presented as mean ± SD, *** *p* < 0.001; Association among FBN2 and d) ATRS and, e) average heal rise, n = 40; f) FBN2 levels among good outcome (GO, n = 20) and poor outcome (PO, n = 20) patients, data presented as mean ± SD, *** *p* < 0.001; g) AUC for eEF2 to report its predictive significance, n = 40; h) Semi-quantitative analysis of eEF2 western blot analysis among good outcome (GO, n = 9) and poor outcome (PO, n = 9) patient samples, data presented as mean ± SD, ** *p* < 0.01, n = 9 per group; i) Positive association between eEF2 and 1-year healing outcome based on western blot analysis, n = 18; j) AUC value of eEF2 based on western blot analysis, n = 18; k) Western blot image of eEF2 and beta-actin (β-actin) in good (G) and poor (P) outcome patients.

To confirm the proteomic data findings for eEF2, western blot analysis was conducted on tissue biopsies collected from same patient cohort used for MS which revealed significantly higher eEF2 levels in good compared to poor outcome patients (Figure 2k). We further studied the ratio of phosphorylated-eEF2 (p-eEF2)/total eEF2 as eEF-2 kinase can modulate the activity of p-eEF2 (Knight et al., 2021). However, no statistical difference was detected among good and poor outcome patients based on both MS and WB data analysis (Supplementary Figure 1). Furthermore, a strong positive association was observed between eEF2 and 1-year clinical outcome (Figure 2i). Together with a strong prognostic significance of the AUC value of 0.9 (Figure 2j), eEF2 was selected as the best prognostic biomarker of Achilles tendon repair and subjected to further mechanistic analyses.

### eEF2 Regulates Collagen Expression During Dense Connective Tissue Repair

To visualize the localization and to confirm eEF2 expression in tissue biopsies, immunohistochemistry (IHC) and immunofluorescence (IF) analysis were performed. Both IHC (Figure 3 a-c) and IF (Figure d-f) analyses clearly demonstrated higher eEF2 expression in the good compared to the poor outcome patients. Further, IHC and IF based semi-quantitative analyses demonstrated that eEF2, which is both a nuclear and cytoplasmic protein (Yao et al., 2014, Sagnol et al., 2014), was mostly located in the ECM area of the tendon tissue (Figure 3 a-b, d-e). To identify how eEF2 improves healing in connective tissues after injury, the association between eEF2 and a classical marker of tendon repair, Collagen type I a1 (Col1a1) was analyzed. Western blotting from protein lysates generated from biopsies used for the MS analysis were used for Col1a1 expression in patients with good and poor outcome. The analysis demonstrated higher Col1a1 expression among patients with better outcome (Figure 3f, g), in accordance with the findings for eEF2 (Figure 1k). The up-regulation of both eEF2 (Figure 1h, k) and Col1a1 was observed in the good compared with the poor outcome group (Figure 2g, h). Interestingly, a strong relationship was observed between eEF2 and Col1a1 (r = 0.61, *p* = 0.007, Figure 3i) levels, suggesting that eEF2 may mediate Col1a1 production to enhance tendon repair.

**Figure 3.**
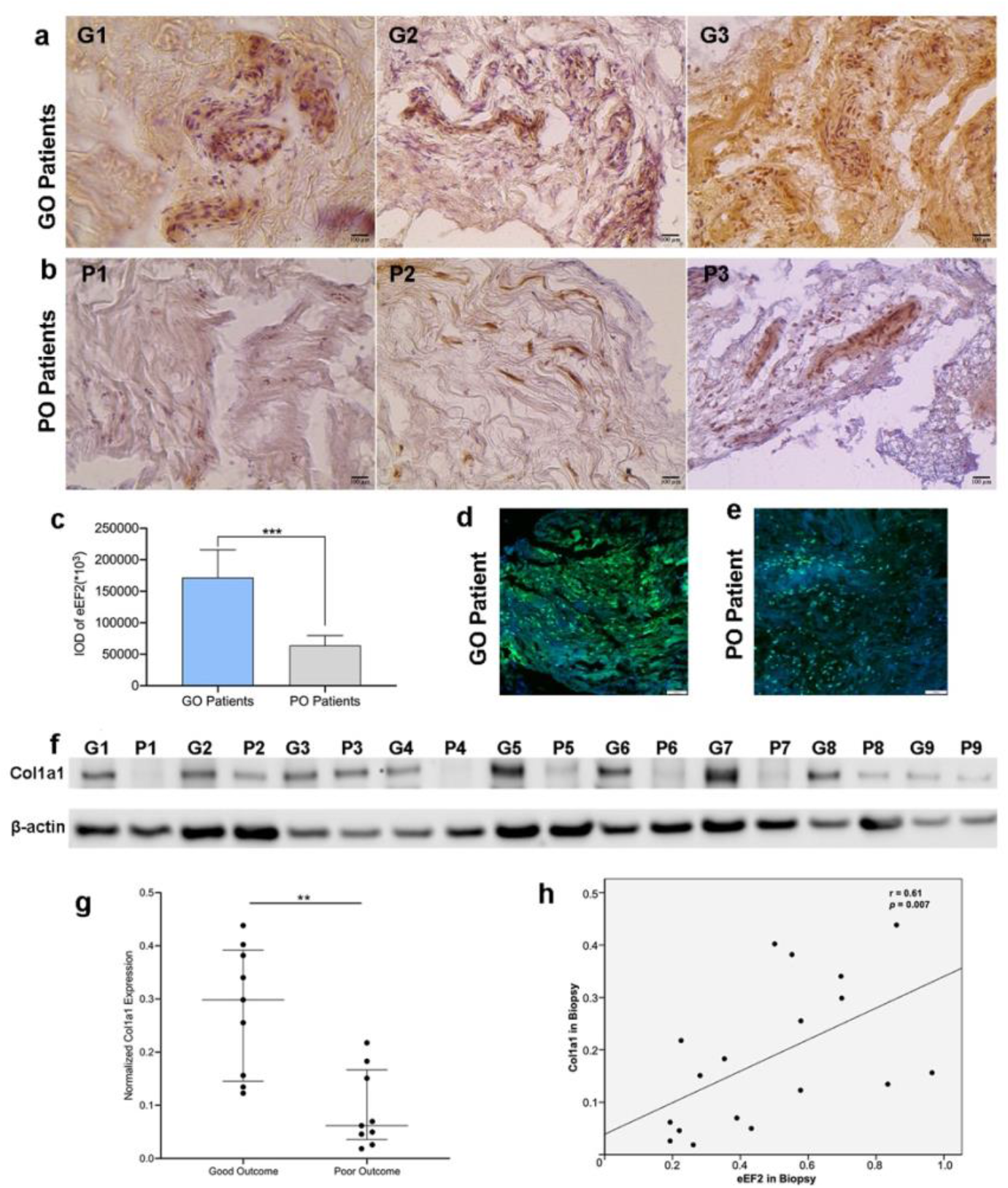
eEF2 expression and localization in tissue biopsies and association with Col1a1. Immunohistochemistry of the eEF2 in a) good outcome (GO) and, b) poor outcome (PO) patients, Scale bars = 100 μm; c) Semi-quantitative analysis of eEF2 expression in the good outcome (GO) and poor outcome (PO) patient samples presented as integrated optical density (IOD), data reported as mean ± SD, *** *p* < 0.001, n = 9 per group. eEF2 immunofluorescence signal in a d) good outcome (GO) and, e) poor outcome (PO) patient sample. Scale bars = 100 μm; f) Western blot analysis and g) semi-quantitative analysis of Col1a1 expression in good and poor outcome patients. Signal intensity was used for quantitative analysis, and the intensity of the house-keeping gene (beta-actin) used for normalization; h) Association among eEF2 and Col1a1 expression, n = 18.

### eEF2 Enhances Dense Connective Tissue Repair during Inflammation by Improving Autophagy

Autophagy, which is a self-renewal mechanism that can degrade and recycle cellular components, plays an essential role in various phases of wound healing (An et al., 2018, Han et al., 2015, Vescarelli et al., 2017). Specifically, in the inflammatory phase autophagy prevents excessive inflammation and can also regulate collagen synthesis (Ren et al., 2022). The bioinformatic analysis, based on the proteomic data also identified that autophagy and collagen expression were improved by eEF2, highlighting a potential role of this novel biomarker during the inflammatory stage of tendon repair (Supplementary Figure 2). Thus, an inflammatory fibroblast injury model was created to confirm and further explore whether eEF2 regulates collagen production by mediating autophagy during the inflammatory phase of repair. An inflammation-induced decline of Col1a1 was detected, and Col1a1 expression was reduced following silencing of eEF2 expression in the cells (Supplementary Figure 3).

Autophagy was induced in the human fibroblast cell line and in primary fibroblasts, leading to an increase in microtubule-associated proteins light chain 3-II (LC3-II) and the ratio of LC3-II/I. The expression of LC3-II and the ratio of LC3-II/I in the fibroblast cell line TERT166 and in primary fibroblasts were significantly reduced when eEF2 was knocked down with si-eEF2, suggesting a positive effect of eEF2 on autophagy (Figure 4a-d). Further exploration of this relationship demonstrated that autophagy led to increases in Col1a1 expression and that this up-regulation was significantly reduced by knocking down eEF2 (Figure 4e-h, i-l). Interestingly, the up- or down-regulation of autophagy and subsequent Col1a1 production resulting from si-eEF2 was approximately equal to the effect induced by an autophagy inhibitor (3-MA) (Figure 4e-l), demonstrating an essential impact of eEF2 on autophagy.

**Figure 4.**
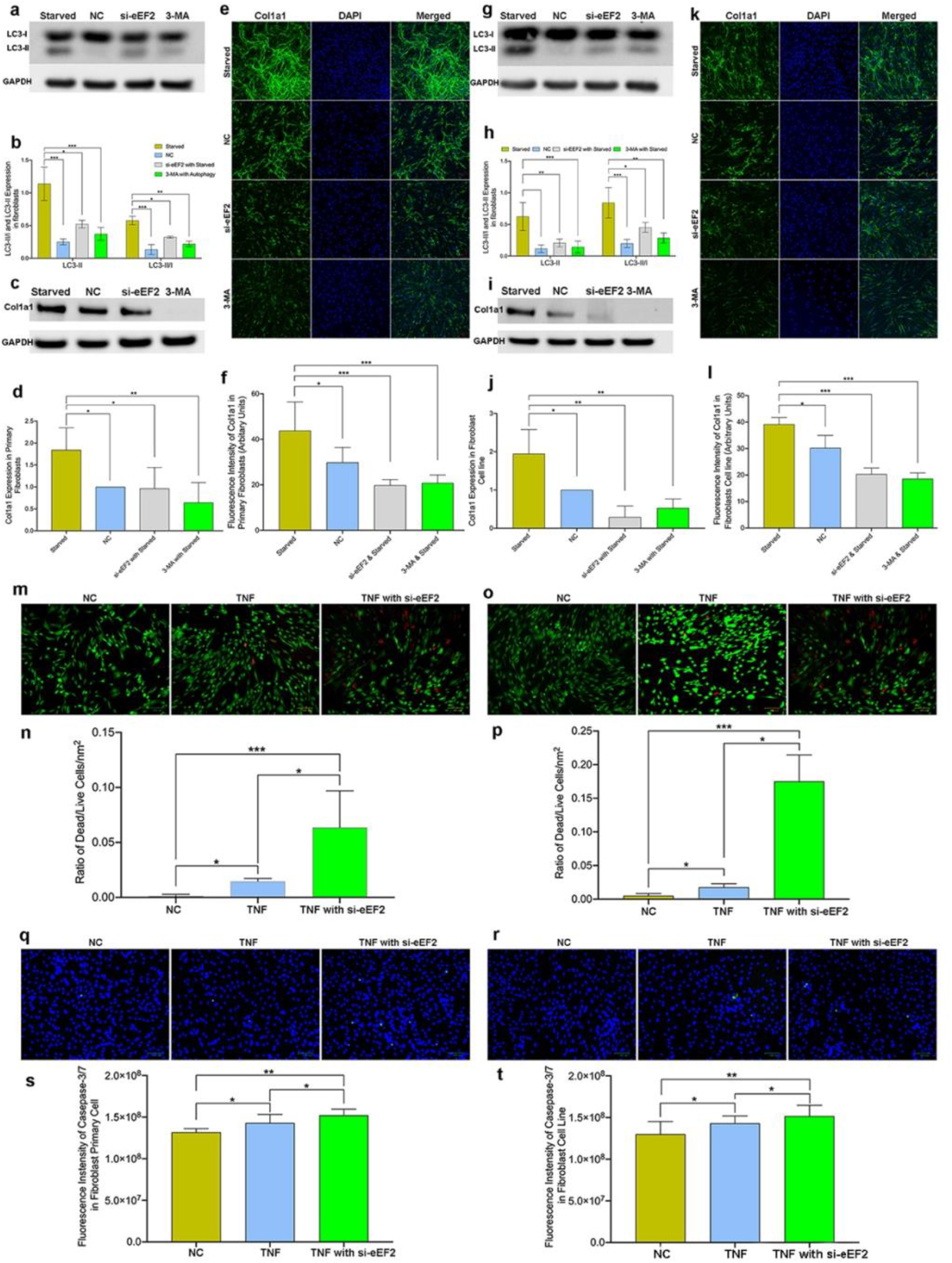
eEF2 enhances healing processes in TNF-induced inflammatory fibroblast models. a-f) eEF2 affects Col1a1 expression through autophagy in primary fibroblasts. a, b) Representative Western blot images and semi-quantitative analysis of LC3-II and LC3-II/LC3-I ratio, c-f) Representative Western blot (c) and confocal images (e) along with semi-quantitative analysis of Col1a1(d, f) in cells treated with normal medium, starved medium for autophagy, si-eEF2 and 3-MA incubation based on autophagy, TNF was used afterward to create an inflammatory model; g-l) eEF2 affects Col1a1 synthesis by autophagy in fibroblast cell line. g, h) Representative Western blot images and semi-quantitative analysis of LC3-II and LC3-II/LC3-I ratio. Representative Western blot (i) and confocal images (k) along with semi-quantitative analysis of Col1a1 (l, j) synthesis in cells treated with normal medium, starved stimulation for autophagy, si-eEF2 and 3-MA incubation based on autophagy; Semi-quantitative analysis demonstrated eEF2 enhanced autophagy and then increased Col1a1 synthesis during inflammation. Signal intensity (a-d, g-j) and fluorescent green intensity (e-f, k-l) were used for semi-quantitative analysis; m-t) Representative images captured by fluorescent microscope demonstrated the cell death and apoptosis when treated with normal condition, TNF and si-eEF2 & TNF: si-eEF2 increases the ratio of dead/live cells among primary fibroblast (m, n) and fibroblast cell line (o, p); si-eEF2 positively associate with cell apoptosis among primary fibroblasts (q, s) and fibroblast cell line (r, t); The ratio of dead/live cells was reported by percentage and the apoptotic level of cells was presented by fluorescent green intensity. Data reported as mean ± SD, * *p* < 0.05, ** *p* < 0.01, *** *p* < 0.001, scale bars = 100 μm, 3 replicates were used for quantitative analysis.

These experimental findings confirmed and expanded the results of the bioinformatic analysis regarding the relationship between eEF2 and autophagy. To our knowledge, this is the first exploration of mechanistic function of eEF2 during regenerative process, highlighting that eEF2 improves Col1a1 production potentially by up-regulating autophagy during inflammation.

### eEF2 Enhances Dense Connective Tissue Repair during Inflammation by Reducing Cell Death and Apoptosis

After tissue injury, the inflammatory stage of healing is comprised of multiple biological processes that are crucial towards tissue repair(Tsuchiya, 2021, Arulselvan et al., 2016, Litwiniuk et al., 2016). Among these processes, we identified that eEF2 not only improved autophagy but also promoted apoptotic processes during tissue repair (Supplementary Figure 2). To confirm the preliminary findings from the bioinformatic and statistical analysis, the ratio of dead/live cells was assessed in two si-eEF2 inflammatory fibroblast models.

The experimental data demonstrated that induction of inflammation led to increase in the ratio of dead/live cells for both primary fibroblasts and the fibroblast cell line (Figure 4m-p). Further, TNF-induced an up-regulation of dead/live ratio that was even higher than when eEF2 was silenced (Figure 4m-p). In addition, the increased ratio of dead/live fibroblasts lead to higher rates of apoptotic cells as observed by caspase-3/7 activity (Figure 4q-t). Hence, this stepwise assessment corroborated that eEF2 decreases cell death and reduces the apoptotic process, highlighting eEF2 as a multi-functional biomarker of tissue repair during the inflammatory stage. Moreover, identical findings from the primary fibroblasts and the fibroblast cell line strengthen the concept that eEF2 plays a vital role by reducing cell death/apoptosis during the inflammatory phase of DCT repair.

### eEF2 Improves Dense Connective Tissue Repair by Enhancing Cell Proliferation

DCTs repair is a complex process in which the inflammatory phase of healing is successively replaced by proliferative healing processes and matrix deposition. Our recent finding based on the proliferative stage of healing (Chen et al., 2022) also detected an increased eEF2 expression in the healing tendons at two weeks post repair surgery. Interestingly, analysis of the MS-based quantitative proteomic data from the early proliferating healing phase; 2-weeks post-surgery, detected elevated levels of eEF2 in the healing compared to intact Achilles tendons (Figure 5a). To further understand the role of eEF2 on proliferative healing processes, the effects of si-eEF2 on proliferating fibroblasts in un-challenged cell line as well in primary fibroblasts were studied.

**Figure 5.**
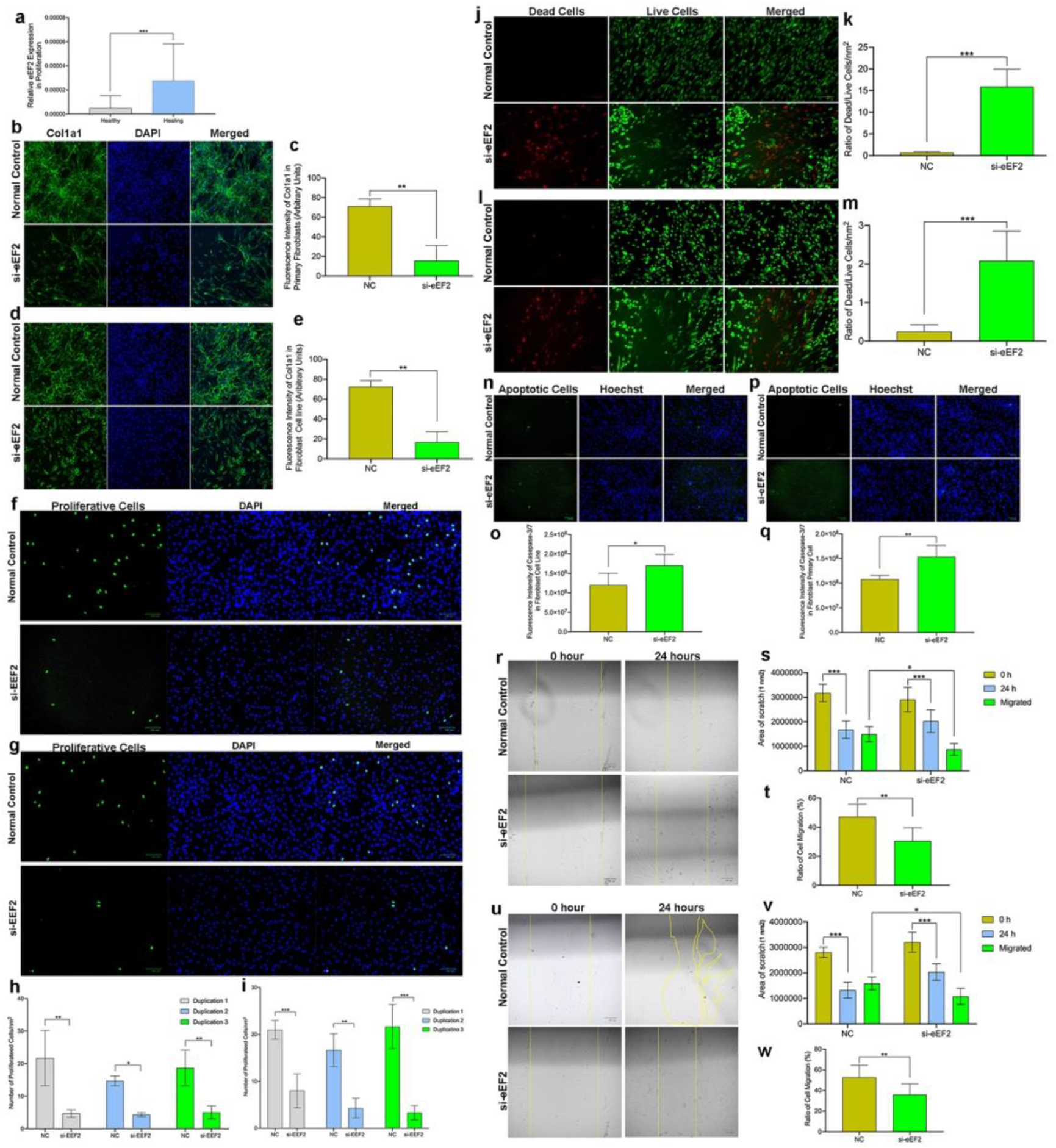
eEF2 enhances fibroblast proliferative processe. a) eEF2 expression from micro-dialysate MS data, micro-dialysate collected from the intact and healing Achilles tendons, 2 weeks post-surgery, *** *p* < 0.001, n = 28 in each group; b-e) Representative confocal images and semi-quantitative analysis of Col1a1 in b,c) primary fibroblasts and, d,e) fibroblast cell line, with and without si-eEF2, data reported as mean ± SD, ** *p* < 0.01, scale bars = 100 μm, n = 3 replicates; f-i) Representative immunofluorescence images and number of proliferating f,h) primary fibroblasts and g,i) fibroblast cell line, with and without si-eEF2. Data reported as mean ± SD, * *p* < 0.05, ** *p* < 0.01, *** *p* < 0.001, scale bars = 100 μm, n =3 replicates; j-m) Representative immunofluorescence images and ratio of dead/live cells from, j,k) primary fibroblasts, and l,m) fibroblast cell line, with and without si-eEF2. Data reported as mean ± SD, *** *p* < 0.001, scale bars = 100 μm, n = 3 replicates; n-q) Representative immunofluorescence images and number of apoptotic cells from n,o) primary fibroblasts, and p,q) fibroblast cell line, with and without si-eEF2. Data reported as mean ± SD, * *p* < 0.05, ** *p* < 0.01, scale bars = 100 μm, n =3 replicates; r-w) Representative images and quantitative analysis of cell migration rate assessed at 0 and 24 hours in r-t) primary fibroblast, and u-w) fibroblast cell line, with and without si-eEF2. Data reported as mean ± SD, * *p* < 0.05, ** *p* < 0.01, *** *p* < 0.001, scale bars = 100 μm, n =3 replicates.

**Figure 6.**
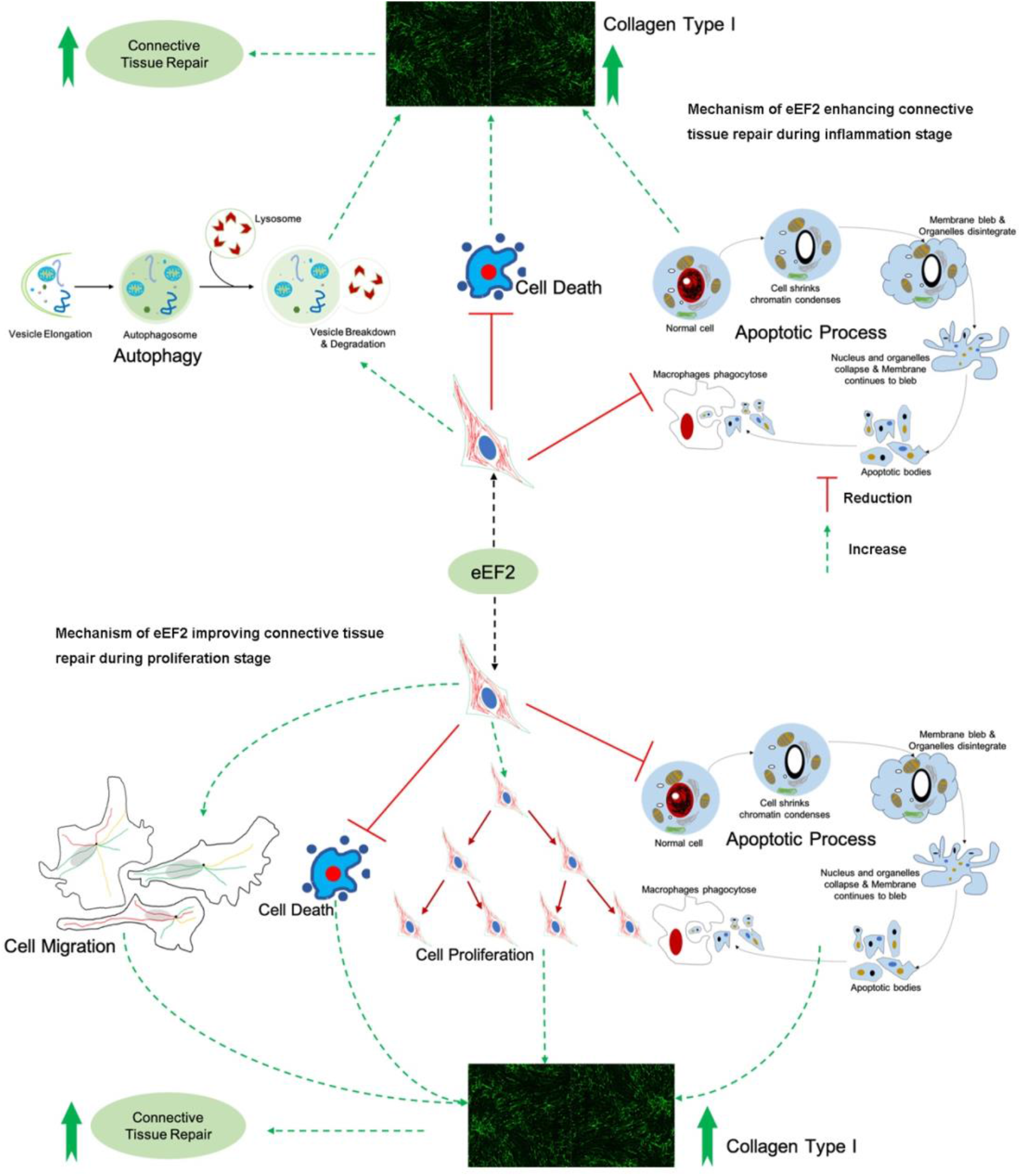
Potential mechanisms and eEF2 mode of action during inflammation and proliferation phases of dense connective tissue repair.

Our first analysis showed a decline in Col1a1 production by knocking down eEF2 (Figure 5b-c, d-e), suggesting an association of eEF2 to this repair-related matrix protein. Quantitative image analysis demonstrated an eEF2-induced cell proliferation among cells treated with si-eEF2 (Figure 5f-i). These observations confirmed and strengthened the preliminary finding, and also extended on the potential role of eEF2 from inflammatory to the early proliferative stages of tendon tissue repair.

### eEF2 Improves Dense Connective Tissue Repair by Reducing Cell Death and Apoptosis during Proliferation

Cell death is an essential yet opposing biological process in relation with proliferation (Guo and Hay, 1999, Oh et al., 2012), while apoptosis is the outcome of cell death. However, the knowledge base for the potentially synchronized regulation of cell apoptotic- and proliferative processes during tissue repair is limited. For the next step of the analysis, the impact of eEF2 on the coordinated cell proliferation and apoptosis leading to connective tissue repair was explored using unchallenged fibroblast models. The experimental observations indicate an opposite effect of eEF2 on cell death when compared with cell proliferation as demonstrated by increases in the ratio of dead/live cells when cells were treated with si-eEF2 (Figure 5j-m). The pathway to cell death was corroborated by demonstrating an increased apoptotic ratio of cells following treatment with si-eEF2 (Figure 5n-q).

Here, these findings for the first-time report that eEF2 induces fibroblast proliferation, and at the same time reduces cell death and apoptosis to improve connective tissue repair and outcomes during an early phase of healing. These observations extended the role of eEF2 at cellular level, but also highlighted its mechanistic function during the early proliferative phase of tissue repair.

### eEF2 Improves Dense Connective Tissue Repair by Enhancing Cell Migration during Proliferation

During wound healing, fibroblast migration to the site of injury is a crucial step to initiate the proliferative healing process (Fronza et al., 2009). Our bioinformatic analysis also observed that eEF2 expression was positively associated with cell migration molecules (Supplementary Figure 4). Thus, we assessed whether eEF2 enhances cell migration during the proliferation phase of tissue healing. For these experiments, *in-vitro* wound models were used in which a scratch was created in monolayer cultured fibroblasts using both the primary cells and the cell line, treated with or without si-eEF2. The findings confirmed the earlier bioinformatic analysis by demonstration of a reduced cell migration area and ratio among cells transfected with si-eEF2 (Figure 5r-w). Quantitative analysis of the results demonstrated a significantly (*p* = 0.01) slower (30%) migration rate in eEF2 knock down cells in comparison with a 47% rate in normal cells when primary cells were used. When the cell line was used in studies, the migration rate was 33% in the si-eEF2 treated cells versus a 55% rate in the control cells. The consistent findings between primary fibroblasts and a fibroblast cell line further support a mechanistic role for eEF2 in cell migration during connective tissue repair.

## Discussion

DCTs are essential to support, protect, and give structure to other tissues and organs in the body. After injury, a reparative process is initiated in dense connective tissues which is highly dependent on protein synthesis. However, biomarkers predictive of good patient outcomes after connective tissue (CT) repair are lacking, particularly those detected during the early phases of the healing process. In this study, the proteomic composition of DCT repair was quantitatively characterized in patients with acute Achilles tendon rupture that were surgically repaired. We successfully generated a comprehensive proteomic file of good versus poor outcome patients. This cohort constitutes a novel collection of human proteomic data related to validated clinical outcomes. These findings obtained from injured Achilles tendon may serve as a representative of more general DCT healing and thus, could greatly improve our understanding of the healing processes involved in DCT repair. We assume that future mining of this proteomic file will expand our understanding of dense connective tissue repair to identify additional potential pathways and potential targets lead to enhancement of healing.

The enrichment analysis of proteomic data identified differentially expressed proteins, which mainly were involved in collagen binding, ECM regulation and organization, and wound healing via cell migration, proliferation and apoptosis. Further exploration of the proteomic file based on a validated clinical database identified a novel biomarker, eEF2 with strong predictability of good healing outcome. By combining bioinformatic analysis along with experimental investigations, the role of eEF2 during tissue repair was explored and verified. Mechanistic exploration of eEF2 identified its potential in vivo role in enhancing autophagy, cell migration and proliferation, as well as reducing cell death and apoptotic processes, all of which contribute to an improved wound healing process.

DCT repair consists of dynamic and overlapping phases, including inflammatory, proliferative and regenerative phases (Qu et al., 2015). To understand the mechanisms of eEF2 in DCT repair we, in a stepwise manner, investigated how this biomarker regulates tissue healing from the inflammatory to the early proliferative healing processes. The finding that eEF2 increases Col1a1 production by positively regulating fibroblast autophagy during inflammation is supported in the literature among different cell types showing an association between autophagy and Col1a1 production (Yang et al., 2020, Vescarelli et al., 2017, Xie et al., 2018, Choi et al., 2020). eEF2 is an elongation factor, which promotes the GTP-dependent translocation of the ribosome, and thus presumably also promotes Col1a1 synthesis in CT repair (Deng et al., 2017, Knight et al., 2021). Col1a1, produced by fibroblasts, encodes the major component of type I collagen, the most abundant collagen in the ECM of healthy connective tissues (Koskinen et al., 2004) (Saraswati et al., 2019, Koos and Bassett, 2018). Moreover, higher expression of type I collagen leads to improved CT repair (Maffulli et al., 2002). Thus, our combined bioinformatics analysis and mechanistic approaches using inflammatory fibroblast models created with both primary human cells and a human cell line, confirmed the hypothesis that eEF2 enhances dense connective tissue repair during the inflammatory phase of healing by increasing autophagy and thereby promoting the production of Col1a1.

The bioinformatic analysis also disclosed a negative relationship between the proteomic profile and apoptosis, which is supported by earlier studies showing that autophagy and apoptosis inhibit each other by protein cleavage or degradation in an enzyme dependent manner (Zhang et al., 2015). In addition, pre-clinical studies have indicated that the inhibition of eEF2 in mice and rat models were found to reduce autophagy, but at the same time activate cell death and apoptosis, as well as decrease protein production in skeletal tissue (Pires Da Silva et al., 2020, Rose et al., 2009). The current findings that eEF2 reduces cell apoptosis as indicated by the down-regulation of the ratio of dead/live cells and apoptotic cells not only extend the role of eEF2 from animal to human species, but also provide mechanistic insight for its role in enhancing DCT repair during the inflammatory phase of healing.

Cell proliferation is the subsequent phase in wound healing (Shah et al., 2012, Landen et al., 2016, Saxena et al., 2019), in which eEF2 has been reported to act as a promoter (Kaleci and Koyuturk, 2020), presumably by improving the cell proliferative process through the PI3K pathway (Shi et al., 2018). However, whether eEF2 plays a key role during the proliferation phase of tissue repair remains unclear. The identification of increased eEF2 levels also at 2-weeks post Achilles tendon surgery (Chen et al., 2022), suggests a role in the early proliferative healing phase as defined in earlier studies of Achilles tendon healing (Ackermann et al., 2013, Alim et al., 2016). Fibroblasts play a vital role during proliferation (Bainbridge, 2013, Tracy et al., 2016, Wong et al., 2007, Addis et al., 2020). Here, we used both unchallenged human primary fibroblasts and a fibroblast cell line, mimicking aspects of the proliferation phase of healing, to investigate the mechanistic role(s) of eEF2 in tissue repair. The combined bioinformatic and mechanistic analysis confirmed that eEF2 enhanced fibroblast proliferation and increased Col1a1 production. In addition, the positive association between eEF2 and Col1a1 extends our understanding of the role of this protein in tissue healing. The coordination between cell proliferative and apoptotic activity also plays a key role in tissue development and homeostasis in the human body (Guo and Hay, 1999). The current findings that eEF2 increases cell proliferation and migration, and at the same time reduces cell death and apoptotic processes, suggest that eEF2 is not only a powerful biomarker, but also provides mechanistic insight of how eEF2 can regulate and enhance DCT repair during the proliferative phase. To the best of our knowledge, this is the first observation of eEF2 as a promoter of multiple phases of tissue repair.

## Conclusion

Taken together, the current findings present eEF2 as a marker prognostic of good quality human Achilles tendon repair, presumably generic to dense connective tissue repair. The bioinformatic and mechanistic analyses confirm that eEF2 promotes healing by regulating processes during the inflammatory as well proliferative phases of tissue repair. The findings presented may lead to the development of targeted treatments which could enhance the long-term healing outcomes for patients suffering with dense connective tissue injuries that currently yield a poor clinical outcome.

## Materials and Methods

### Ethical Approval and Consent to Participate

This study was conducted with the approval from the Regional Ethical Review Committee in Sweden and followed all guidelines according to the Declaration of Helsinki. Written consent was obtained from all patients.

### Patient Eligibility Criteria and Randomization

This was a retrospective study of 40 patients suffering from an acute Achilles tendon rupture (ATR) who underwent reconstruction surgery at the Karolinska University Hospital. The majority of the ATR-cases (90%) were sports-related. During the surgery, tendon biopsies were taken from the ruptured area and stored at minus 80°C for future analysis. Patients were randomly selected from a patient cohort who participated in randomized control trials at the hospital. All the participants received the same anesthetic and surgical procedure to repair mid-substance tears within 2-7 days of injury and followed exclusion criteria as reported previously (Chen et al., 2021, Alim et al., 2016).

### Patient-reported and Functional Outcome

The patient-reported outcomes were evaluated 1-year postoperatively using validated questionnaires: Achilles Tendon Total Rupture Score (ATRS), Foot and Ankle Outcome Score (FAOS) and functional outcome was measured using the Heel-rise Test (HRT). All the methodology, rules and regulations of patient outcome measurement followed the same description as in our previous studies (Svedman et al., 2018, Alim et al., 2016, Abdul Alim et al., 2018, Chen et al., 2021).

### Mass Spectrometry (MS)

#### Protein Extraction and Digestion of Tissue Samples

Pulverized patient tissue samples were solubilized in 8M urea and 100 mM NaCl with 1% ProteaseMAX (Promega) in 100 mM ammonium bicarbonate (AmBic) and mixed vigorously. Low binding silica beads (400 μm, Ops Diagnostics, Lebanon NJ) were added to each sample and vortexed at high speed. Subsequently, samples were quickly frozen and subsequently then thawed and subjected twice to disruption of the tissue on a Vortex Genie disruptor for 2 min before addition of AmBic, urea and NaCl. Following centrifugation, the supernatant was transferred to new tubes and 50 mM AmBic was added and the mixture was vortexed vigorously.

Proteins were reduced with 100 mM dithiothreitol in 50 mM AmBic, incubated at 37°C and alkylated with 100 mM iodoacetamide in 50 mM AmBic in the dark. Proteolytic digestion was performed overnight. The reaction was stopped with concentrated formic acid and the samples were then cleaned on a C18 Hypersep plate (bed volume of 40 μL, Thermo Scientific) and dried in a vacuum concentrator (miVac, Thermo Scientific).

#### RPLC-MS/MS Analysis

Briefly, reversed phase liquid chromatographic separations of peptides were performed on a C18 EASY-spray and C18 trap columns connected to an Ultimate 3000 UPLC system (Thermo Scientific). Mass spectra were acquired on an Q Exactive HF mass spectrometer (Thermo Scientific), targeting 5×10^6^ ions with maximum injection time of 100 ms, followed by data-dependent higher-energy collisional dissociation (HCD) fragmentations from precursor ions with a charge state.

#### Proteomic Data Analysis, Protein Identification and Quantification

Raw files were imported to Proteome Discoverer v2.3 (Thermo Scientific) and analyzed using the SwissProt protein database with the Mascot v 2.5.1 (MatrixScience Ltd., UK) search engine. MS/MS spectra were matched with The Human Uniport database (last modified: 3 September 2020; ID: UP000005640; 75,777 proteins) using the MSFragger database engine (Kong et al., 2017).

Protein abundance was calculated based on normalized spectrum intensity (LFQ intensity), and an intensity-based absolute quantification (iBAQ) algorithm was used for normalization(Schwanhausser et al., 2011). A *p* value < 0.05 and fold change (FC) > 2 were set as significant threshold to identify differentially expressed proteins between good and poor outcome patients. Gene Ontology (GO) annotation and KEGG database were used to identify the biological functions of enriched proteins. Enrichment analysis was performed with GO annotations on STRING v11.0 (http://string-db.org) database for the confirmation of protein-protein interaction network among subgroups. Gene set enrichment analysis (GSEA) and single-sample GSEA (ssGSEA), based on the GO and KEGG databases were used to detect potential pathways involved in the candidate identified biomarkers.

#### Cell Culture

The use of fibroblast cells for connective tissue studies has been previously reported (Chu et al., 2020). Human primary fibroblasts (#C-12302, Promo Cells) and the fHDF/TERT166 human foreskin fibroblast cell line (#CHT-031-0166, Evercyte, Austria) were used for the *in vitro* studies and for confirmation of the in vivo findings. Cells were cultured in Dulbecco’s Modified Eagle Medium/F12 (DMEM/F12, Gibco) supplemented with 10% fetal bovine serum (FBS, Gibco) and 1% penicillin-streptomycin (PEST) (Gibco) at 37°C in a humidified atmosphere containing 5% CO_2_. After reaching 80-90% confluency, cells were dissociated with 0.25% trypsin-ethylenediaminetetraacetic acid (Trypsin-EDTA) (Thermo Fisher Scientific) and seeded in 12-well plates at 2.5 × 10^5^ cells/well supplemented with DMEM/F-12 medium for further experiments.

#### Cell Transfection

Human fibroblasts were seeded and transfected with eEF2 silencing RNA (si-eEF2) (#4392420, Ambion) to detect the potential role of eEF2 in tendon healing. Cells were stimulated with 10 ng/ml TNF for 24 hours before being transfected with si-eEF2. Cells exposed to silencing RNA for eEF2 and negative controls were transfected for 48 hours at a final concentration of 100 nM in each well using Lipofectamine™ 3000 transfection reagent (Thermo Fisher Scientific) and diluted with Opti-MEM reagent.

#### Inflammatory Fibroblast Injury Model

Recombinant human tumor necrosis factor alpha (TNF-α) (#300-01A, PeproTech) was used to establish an inflammatory insult to the fibroblasts after transfection. Bovine serum albumin (BSA), 0.2% was used for dissolving the TNF, and 10 ng/ml was used for fibroblast stimulation (Lee et al., 2021).

#### Immunohistochemistry (IHC) and Immunofluorescence (IF)

Human tendon biopsies were cut into 7 μm sections using a cryostat. Briefly, tissue sections were incubated with 2% H_2_O_2_ and blocked with 5% bovine serum albumin (BSA) at room temperature. Monoclonal antibody to eEF2 (#MA5-42937; Thermo Fisher Scientific) and a secondary antibody, goat anti-rabbit (#31460; Thermo Fisher Scientific, 1:250) was used for immunolocalization. The bound antibody was visualized with diaminobenzidine (DAB, Vector Laboratories, Inc. Burlingame, CA, USA) and counterstained with Hematoxylin (Vector Laboratories, Inc. Burlingame, CA, USA). This step was followed by dehydration in alcohol and xylene before mounting with pertex (Sigma). To demonstrate specificity of staining, primary antiserum was either omitted or replaced by normal rabbit IgG.

For fluorescence staining, slides were blocked with 2% BSA in 1 mol/L phosphate buffered saline (PBS) and subsequently incubated with the primary antibody against eEF2 overnight at 4°C and then incubated with Alexa Fluor 488-labeled Goat anti-rabbit IgG secondary antibody at room temperature. All the slides were stained with antifade reagent (with DAPI) and mounted. Negative controls were prepared by omitting the primary antibody or including a rabbit IgG1 (M5284, 1:100 dilution) isotype control. Data analysis was performed using ImageJ software.

Cells were plated on the glass chamber slides and treated with starved stimulation for autophagy, eEF2 siRNA and 3-Methyladenie (3-MA), respectively. Next, the cells were washed, fixed with 4% paraformaldehyde (PFA), blocked with 5% BSA in PBS, permeabilized with 0.1% Triton X-100, incubated with anti-Col1a1 antibody (CST) and followed by Alexa-Fluor-488 conjugated secondary antibody (#A-11008; Thermo Fisher Scientific). All the slides were stained with the antifade reagent (with DAPI) and mounted. The single-stained and merged images were performed using a Zeiss LSM 880 confocal microscopy (without AiryScan, CMM Karolinska Institutet), and the data was analyzed using ImageJ software.

#### Cell Proliferation Assay

A Click-iT ^®^ EdU AlexaFluor 488 ^®^ Imaging Kit (Thermo Fisher Scientific) was used to identify the changes in proliferation rates of human fibroblast primary cells and the cell line. Normal, transfected and 3-MA stimulated cells were incubated with EdU media for 2 hours before fixation with 4% PFA. Each section was incubated with freshly prepared Click-iT^®^ reaction cocktail as instructed in the kit. BSA (3%) in PBS was used to wash the slides twice for 5 min. Cell nuclei were stained with Hoechst 33342 for 5 min and the slides were subsequently washed with PBS for 2 × 5 min. All the steps of the EdU assay were performed while avoiding light and the slides were never allowed to dry out.

#### Cell Apoptosis Assay

Live/Dead™ Cell Imaging Kit and CellEvent™ Caspase-3/7 Green Detection Reagent (Thermo Fisher Scientific) were used for detection of cell apoptosis based on the manufacturer instructions. Cells were treated with si-eEF2, fixed with 4% PFA (specifically for casepase-3/7) and stained using an annexin V-FITC, PI and Caspase-3/7 green detection reagent in dark. Hoechst 33342 was used for cell nucleus staining. Finally, cells were assessed and quantified using the fluorescence microscope. Image J and SoftMax Pro 7.0.3 were used to perform the data analysis.

#### Scratch Wound Assay

A cell scratch assay was used to explore the role of eEF-2 on cell migration. Based on an unchallenged fibroblast model, cells were seeded in 6-well plates and a scratch was created using a pipette tip on the confluent monolayer cell culture. Next, cells were transfected with si-eEF2 and compared with a control group, all the subgroups were treated with no-FBS to block cell proliferation. After 24 hours, light microscopy and ImageJ were used to assess the results.

#### Western Blot Analysis

Briefly, 5 μg of protein per well was loaded, separated by gel electrophoresis (4-12% Bis-Tris, Invitrogen), and transferred to nitrocellulose membranes. The membranes were incubated with 5% nonfat milk in 1x tris-buffered saline with 0.1% tween (TBST) to block unspecific binding sites, and the membranes were subsequently covered with different primary antibodies overnight at 4°C (eEF2 0.1μg /ml, Col1a1 0.1μg /ml, β-actin 0.02 μg /ml, GAPDH 0.01μg /ml, Cell Signaling). After washing the membranes, they were incubated with secondary antibody conjugated to HRP (anti-rabbit IgG and anti-mouse IgG; Cell Signaling). The chemiluminescence signal was initiated by using SuperSignal West Pico PLUS kit (Thermo Fisher Scientific). Chemiluminescence was identified and quantified by using biorad ChemiDoc MP Imaging and analysis was performed in Image Lab. Immunopositive bands were normalized with the bands of GAPDH or β-actin and final results were presented as fold change from control values.

#### Statistical analysis

Statistical analysis and data plotting were performed with SPSS (IBM SPSS, v26.0), GraphPad Prism 8.0 and R. Skewness was checked for all the variables. Standard descriptive statistics were used to summarize clinical variables such as mean and SD. Differential comparison of protein expressions was calculated using the unpaired Mann-Whitney *U* test, a *p* value < 0.05 and FC > 2 were considered as statistically significant. The patient outcome measurements were associated with the corresponding proteomic files by means of univariate analysis. Outcome measures that were significant in univariate analysis were subsequently used as dependent variables in a multiple-linear regression model with independent variables, and an adjusted *p* value < 0.05 was set as the significance threshold.

## Reporting summary

Further information on research design is available in the Nature Research Reporting Summary linked to this article.

## Acknowledgments

We would like to thank LB for the collection of human samples. Dr. AV at the Proteomics Biomedicum, Karolinska Institutet, Sweden for excellent help in MS data analysis and interpretation.

## Declarations

### Ethical approval and consent to participate

This study was conducted after approval from the Regional Ethical Review Committee in Sweden (Reference no. 2009/2079-31/2: 2013/1791-31/3) and followed all guidelines according to the Declaration of Helsinki.

### Trial registration

NCT02318472 registered 17 December 2014 and NCT01317160 registered 17 March 2011, with URL http://clinicaltrials.gov/ct2/show/NCT02318472 and http://clinicaltrials.gov/ct2/show/study/NCT01317160.

### Consent for publication

All the co-authors give their consent for the findings related to this manuscript to be published.

### Availability of data and materials

MS proteomic data files are deposited at the ProteomeXchange Consortium (http://proteomecentral.proteomexchange.org) through the PRIDE partner repository (https://www.ebi.ac.uk/pride/) under the dataset PXD029202 and PXD033163. The extracted protein LFQ intensity and GSEA data are provided as supplementary files. Clinical data was collected from the database at the section of Orthopedics, Department of Molecular Medicine and Surgery, Karolinska Institutet.

### Author contribution

The project organization, training, resources and funding acquisition by PW. Ackermann and AS. Ahmed; Data analysis by J. Wang and X. Wu; Data curation by N. Simon, XJ. Wu and J. Chen; Visualization by J. Wang, DA. Hart and AS. Ahmed; Experiment completed by J. Chen; Writing original draft by J. Chen; Manuscript reviewing and editing by N. Simon, CI. Svensson, DA. Hart, AS. Ahmed and PW. Ackermann.

---

Original Western blot image of Col1a1 in human tissue, n=18

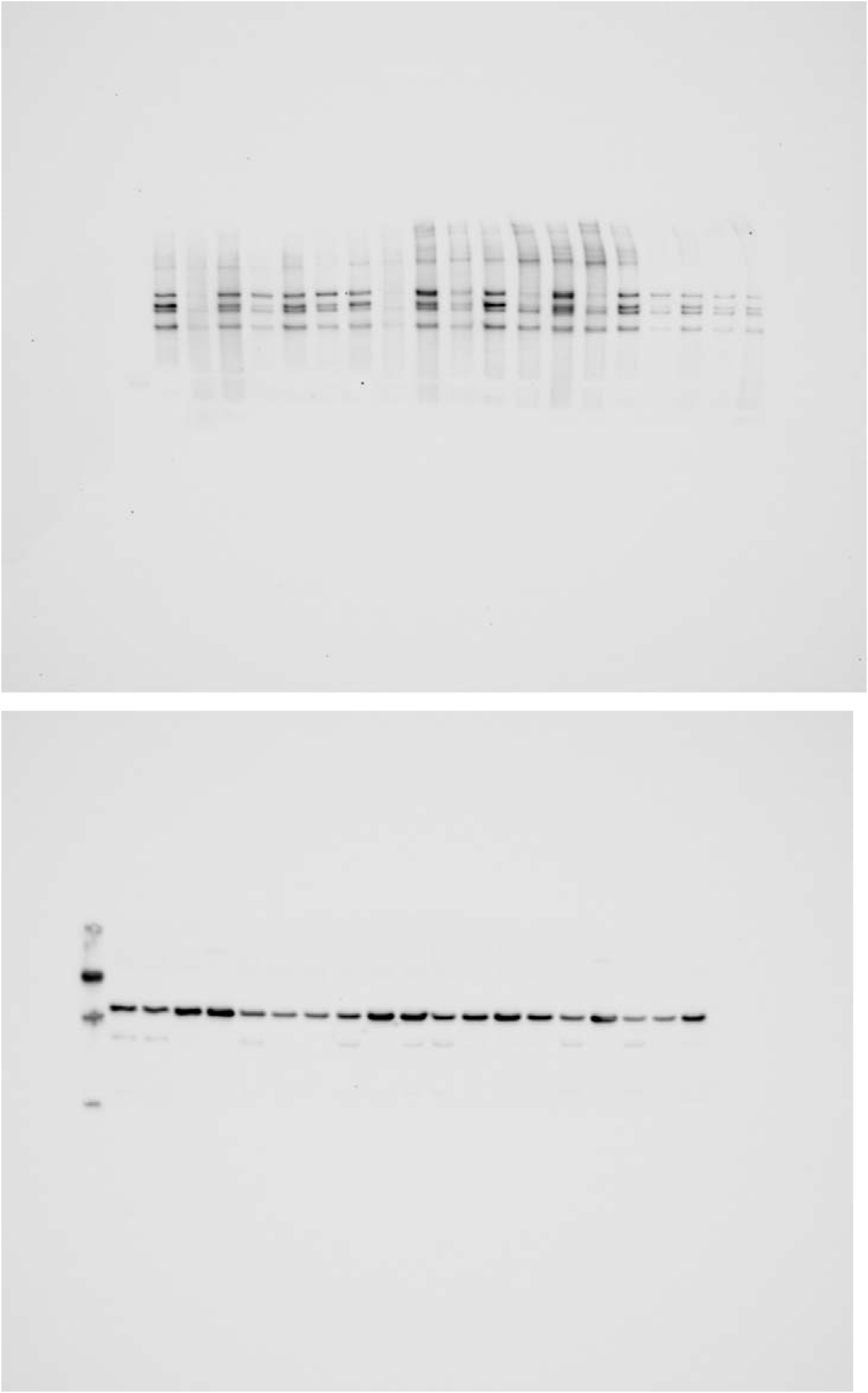

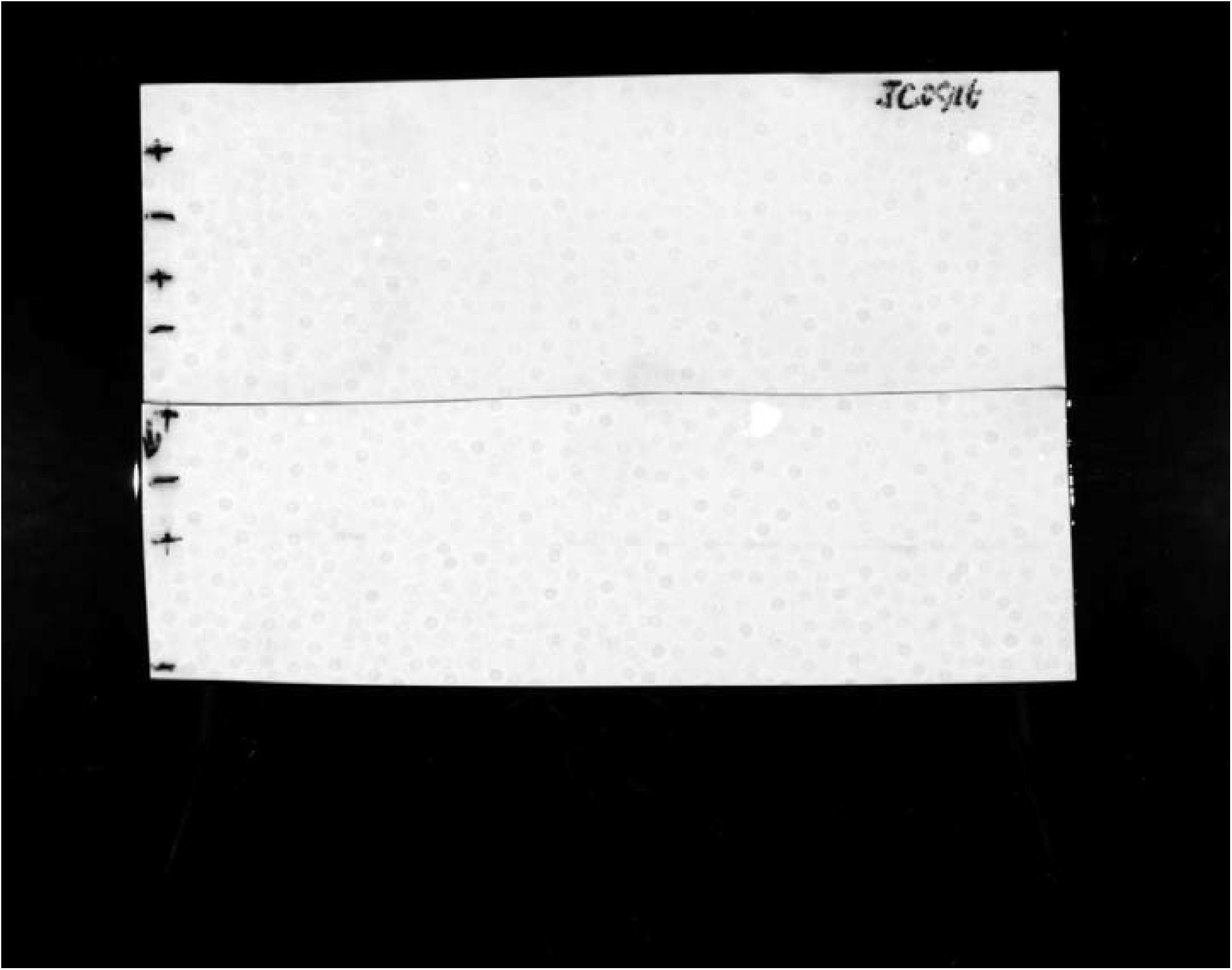

Original Western blot image of Col1a1 (220kDa) in primary fibroblast and fibroblast cell line:

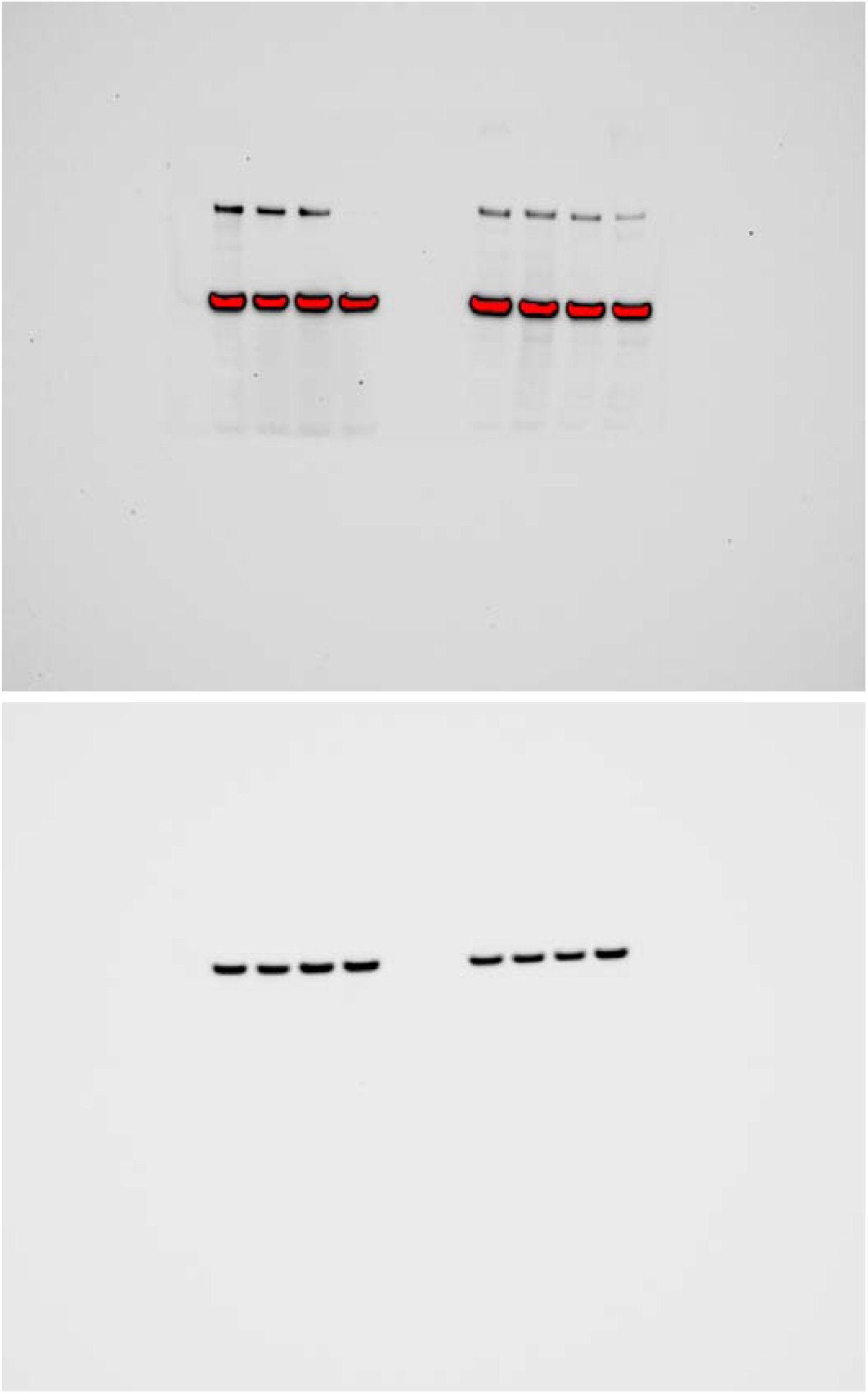

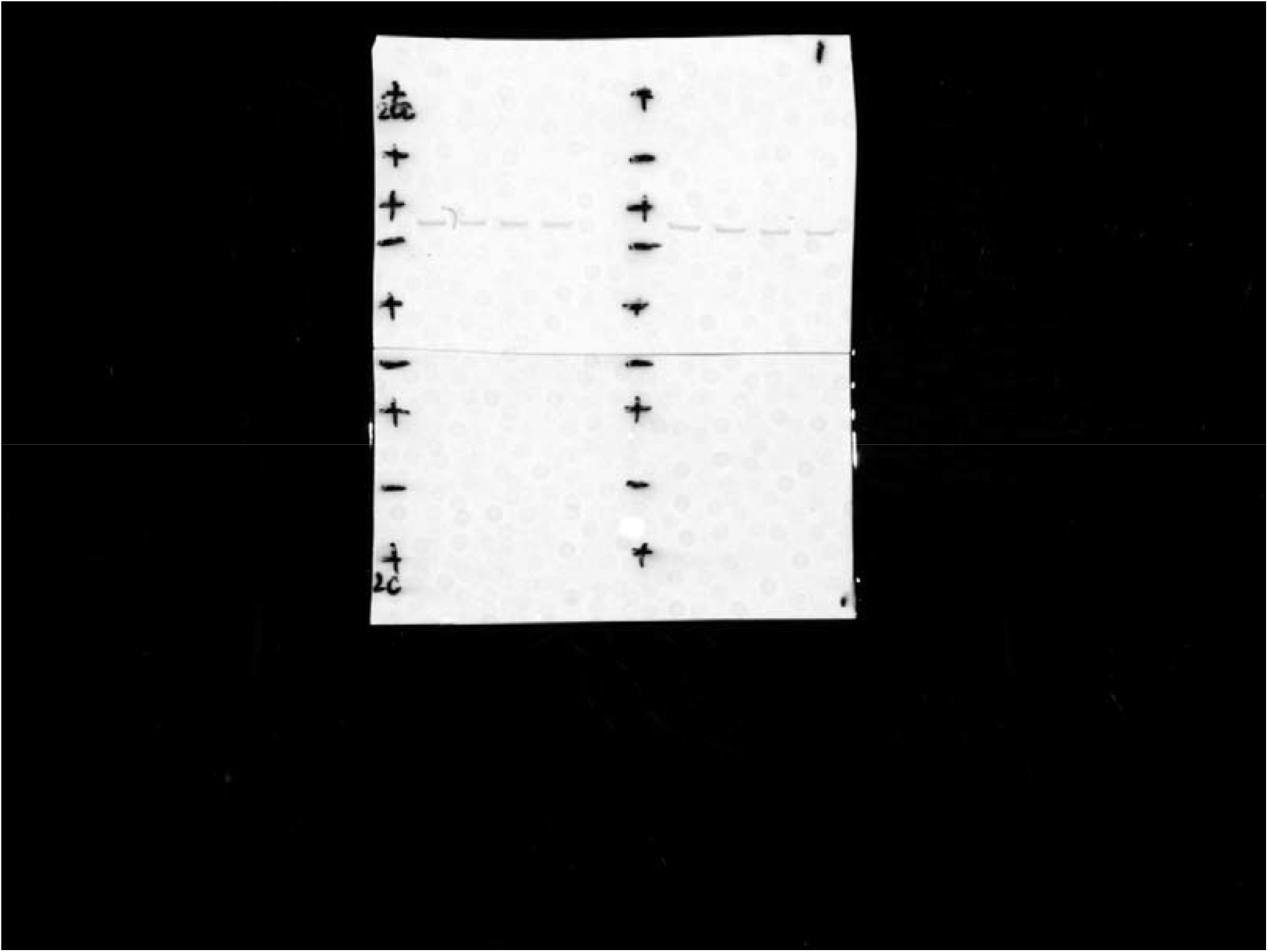

Original Western blot image of eef2 in human tissues, n=18

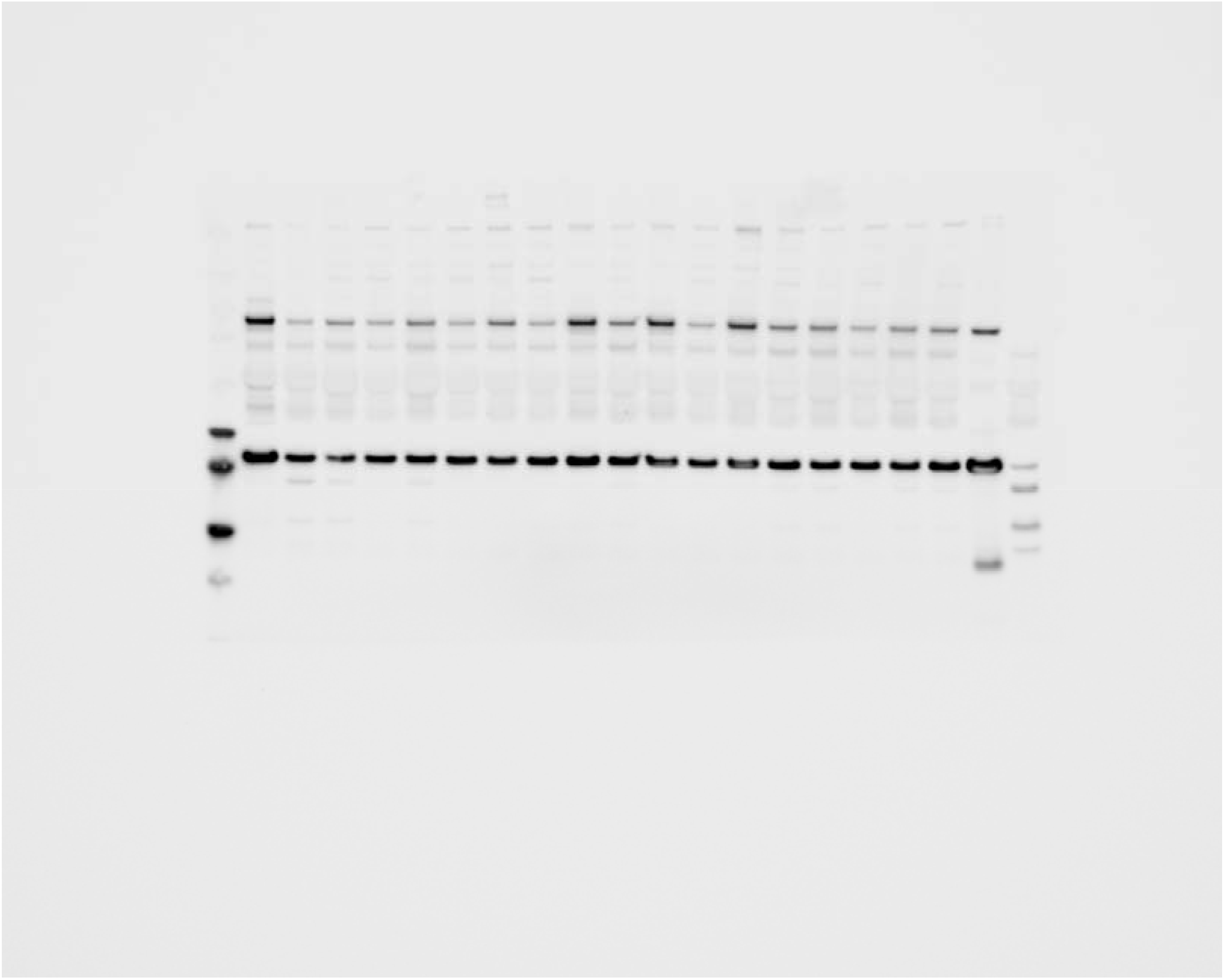

Original Western blot image of LC3-I and II in primary fibroblast and fibroblast cell line:

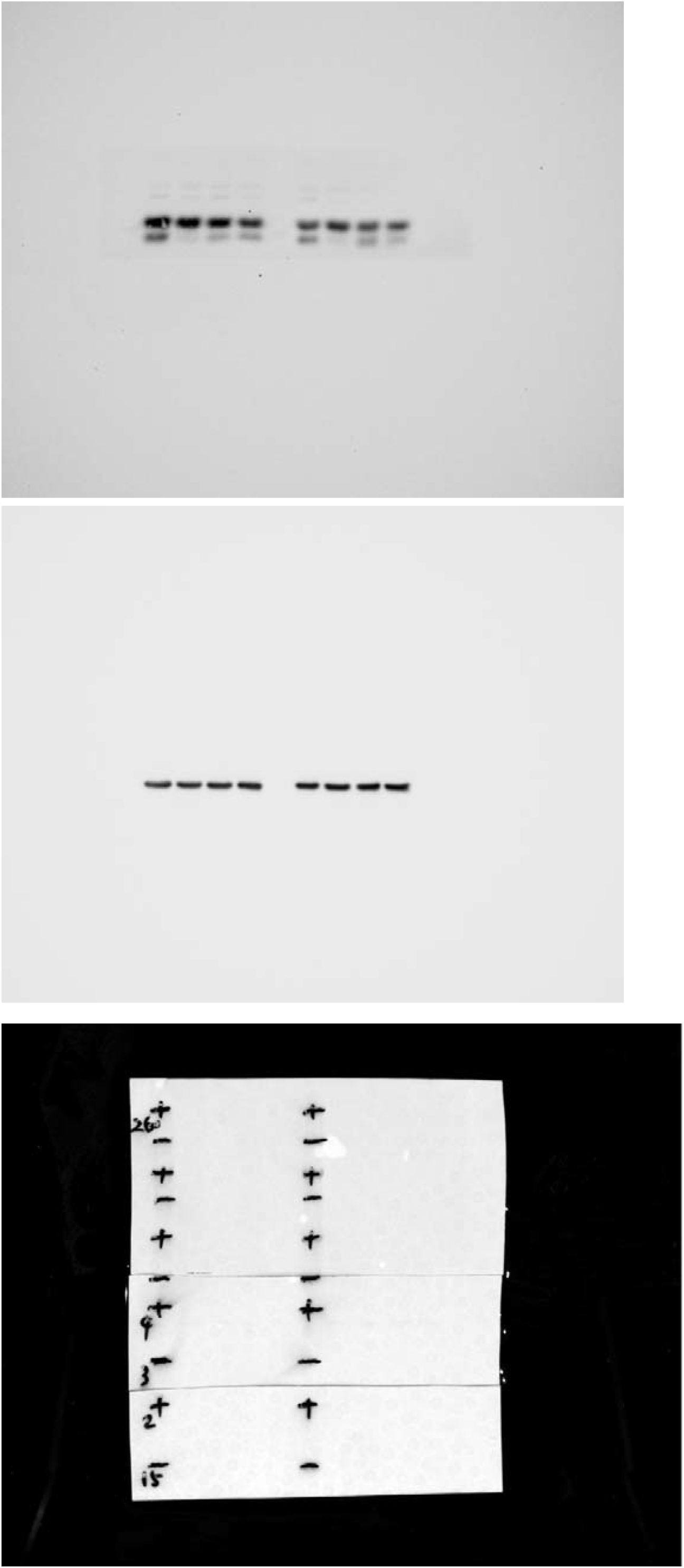

## Reference

Abdul Alim, M., Domeij-Arverud, E., Nilsson, G., Edman, G. & Ackermann, P. W. 2018. Achilles tendon rupture healing is enhanced by intermittent pneumatic compression upregulating collagen type I synthesis. Knee Surg Sports Traumatol Arthrosc, 26, 2021–2029.

Ackermann, P. W., Domeij-Arverud, E., Leclerc, P., Amoudrouz, P. & Nader, G. A. 2013. Anti-inflammatory cytokine profile in early human tendon repair. Knee Surg Sports Traumatol Arthrosc, 21, 1801–6.

Addis, R., Cruciani, S., Santaniello, S., Bellu, E., Sarais, G., Ventura, C., Maioli, M. & Pintore, G. 2020. Fibroblast Proliferation and Migration in Wound Healing by Phytochemicals: Evidence for a Novel Synergic Outcome. Int J Med Sci, 17, 1030–1042.

Alim, M. A., Svedman, S., Edman, G. & Ackermann, P. W. 2016. Procollagen markers in microdialysate can predict patient outcome after Achilles tendon rupture. BMJ Open Sport Exerc Med, 2, e000114.

An, Y., Liu, W. J., Xue, P., Ma, Y., Zhang, L. Q., Zhu, B., Qi, M., Li, L. Y., Zhang, Y. J., Wang, Q. T. & Jin, Y. 2018. Autophagy promotes MSC-mediated vascularization in cutaneous wound healing via regulation of VEGF secretion. Cell Death Dis, 9, 58.

Anderson-Baucum, E., Pineros, A. R., Kulkarni, A., Webb-Robertson, B. J., Maier, B., Anderson, R. M., Wu, W., Tersey, S. A., Mastracci, T. L., Casimiro, I., Scheuner, D., Metz, T. O., Nakayasu, E. S., Evans-Molina, C. & Mirmira, R. G. 2021. Deoxyhypusine synthase promotes a pro-inflammatory macrophage phenotype. Cell Metab, 33, 1883–1893 e7.

Arulselvan, P., Fard, M. T., Tan, W. S., Gothai, S., Fakurazi, S., Norhaizan, M. E. & Kumar, S. S. 2016. Role of Antioxidants and Natural Products in Inflammation. Oxid Med Cell Longev, 2016, 5276130.

Bainbridge, P. 2013. Wound healing and the role of fibroblasts. J Wound Care, 22, 407-8, 410-12.

Baranzini, N., Pulze, L., Tettamanti, G., Acquati, F. & Grimaldi, A. 2021. HvRNASET2 Regulate Connective Tissue and Collagen I Remodeling During Wound Healing Process. Front Physiol, 12, 632506.

Chen, J., Svensson, J., Sundberg, C. J., Ahmed, A. S. & Ackermann, P. W. 2021. FGF gene expression in injured tendons as a prognostic biomarker of 1-year patient outcome after Achilles tendon repair. J Exp Orthop, 8, 20.

Chen, J., Wang, J., Hart, D. A., Ahmed, A. S. & Ackermann, P. W. 2022. Complement factor D as a predictor of Achilles tendon healing and long-term patient outcomes. FASEB J, 36, e22365.

Choi, H., Simpson, D., Wang, D., Prescott, M., Pitsillides, A. A., Dudhia, J., Clegg, P. D., Ping, P. P. & Thorpe, C. T. 2020. Heterogeneity of proteome dynamics between connective tissue phases of adult tendon. Elife, 9.

Chu, J., Lu, M., Pfeifer, C. G., Alt, V. & Docheva, D. 2020. Rebuilding Tendons: A Concise Review on the Potential of Dermal Fibroblasts. Cells, 9.

Deng, Z., Luo, P., Lai, W., Song, T., Peng, J. & Wei, H. K. 2017. Myostatin inhibits eEF2K-eEF2 by regulating AMPK to suppress protein synthesis. Biochem Biophys Res Commun, 494, 278–284.

Fischer, A., Wannemacher, J., Christ, S., Koopmans, T., Kadri, S., Zhao, J., Gouda, M., Ye, H., Muck-Hausl, M., Krenn, P. W., Machens, H. G., Fassler, R., Neumann, P. A., Hauck, S. M. & Rinkevich, Y. 2022. Neutrophils direct preexisting matrix to initiate repair in damaged tissues. Nat Immunol, 23, 518–531.

Fronza, M., Heinzmann, B., Hamburger, M., Laufer, S. & Merfort, I. 2009. Determination of the wound healing effect of Calendula extracts using the scratch assay with 3T3 fibroblasts. J Ethnopharmacol, 126, 463–7.

Grol, M. W., Haelterman, N. A., Lim, J., Munivez, E. M., Archer, M., Hudson, D. M., Tufa, S. F., Keene, D. R., Lei, K., Park, D., Kuzawa, C. D., Ambrose, C. G., Eyre, D. R. & Lee, B. H. 2021. Tendon and motor phenotypes in the Crtap(-/-) mouse model of recessive osteogenesis imperfecta. Elife, 10.

Guo, M. & Hay, B. A. 1999. Cell proliferation and apoptosis. Curr Opin Cell Biol, 11, 745–52.

Han, Y. F., Sun, T. J., Han, Y. Q., Xu, G., Liu, J. & Tao, R. 2015. Clinical perspectives on mesenchymal stem cells promoting wound healing in diabetes mellitus patients by inducing autophagy. Eur Rev Med Pharmacol Sci, 19, 2666–70.

Hussein, A. I., Mancini, C., Lybrand, K. E., Cooke, M. E., Matheny, H. E., Hogue, B. L., Tornetta, P., 3RD & Gerstenfeld, L. C. 2018. Serum proteomic assessment of the progression of fracture healing. J Orthop Res, 36, 1153–1163.

Kaleci, B. & Koyuturk, M. 2020. Efficacy of resveratrol in the wound healing process by reducing oxidative stress and promoting fibroblast cell proliferation and migration. Dermatol Ther, 33, e14357.

Kim, M. S., Pinto, S. M., Getnet, D., Nirujogi, R. S., Manda, S. S., Chaerkady, R., Madugundu, A. K., Kelkar, D. S., Isserlin, R., Jain, S., Thomas, J. K., Muthusamy, B., Leal-Rojas, P., Kumar, P., Sahasrabuddhe, N. A., Balakrishnan, L., Advani, J., George, B., Renuse, S., Selvan, L. D., Patil, A. H., Nanjappa, V., Radhakrishnan, A., Prasad, S., Subbannayya, T., Raju, R., Kumar, M., Sreenivasamurthy, S. K., Marimuthu, A., Sathe, G. J., Chavan, S., Datta, K. K., Subbannayya, Y., Sahu, A., Yelamanchi, S. D., Jayaram, S., Rajagopalan, P., Sharma, J., Murthy, K. R., Syed, N., Goel, R., Khan, A. A., Ahmad, S., Dey, G., Mudgal, K., Chatterjee, A., Huang, T. C., Zhong, J., Wu, X., Shaw, P. G., Freed, D., Zahari, M. S., Mukherjee, K. K., Shankar, S., Mahadevan, A., Lam, H., Mitchell, C. J., Shankar, S. K., Satishchandra, P., Schroeder, J. T., Sirdeshmukh, R., Maitra, A., Leach, S. D., Drake, C. G., Halushka, M. K., Prasad, T. S., Hruban, R. H., Kerr, C. L., Bader, G. D., Iacobuzio-Donahue, C. A., Gowda, H. & Pandey, A. 2014. A draft map of the human proteome. Nature, 509, 575–81.

Knight, J. R., Vlahov, N., Gay, D. M., Ridgway, R. A., Faller, W. J., Proud, C., Mallucci, G. R., Von Der Haar, T., Smales, C. M., Willis, A. E. & Sansom, O. J. 2021. Rpl24(Bst) mutation suppresses colorectal cancer by promoting eEF2 phosphorylation via eEF2K. Elife, 10.

Kong, A. T., Leprevost, F. V., Avtonomov, D. M., Mellacheruvu, D. & Nesvizhskii, A. I. 2017. MSFragger: ultrafast and comprehensive peptide identification in mass spectrometry-based proteomics. Nat Methods, 14, 513–520.

Koos, J. A. & Bassett, A. 2018. Genetics Home Reference: A Review. Med Ref Serv Q, 37, 292–299.

Koskinen, S. O. A., Heinemeier, K. M., Olesen, J. L., Langberg, H. & Kjaer, M. 2004. Physical exercise can influence local levels of matrix metalloproteinases and their inhibitors in tendon-related connective tissue. Journal of Applied Physiology, 96, 861–864.

Landen, N. X., Li, D. & Stahle, M. 2016. Transition from inflammation to proliferation: a critical step during wound healing. Cell Mol Life Sci, 73, 3861–85.

Lee, J. M., Hwang, J. W., Kim, M. J., Jung, S. Y., Kim, K. S., Ahn, E. H., Min, K. & Choi, Y. S. 2021. Mitochondrial Transplantation Modulates Inflammation and Apoptosis, Alleviating Tendinopathy Both In Vivo and In Vitro. Antioxidants (Basel), 10.

Li, B. & Wang, J. H. 2011. Fibroblasts and myofibroblasts in wound healing: force generation and measurement. J Tissue Viability, 20, 108–20.

Litwiniuk, M., Krejner, A., Speyrer, M. S., Gauto, A. R. & Grzela, T. 2016. Hyaluronic Acid in Inflammation and Tissue Regeneration. Wounds, 28, 78–88.

Liu, J. Y., Luo, C. Q., Yin, Z. Q., Li, P., Wang, S. H., Chen, J., He, Q. Y. & Zhou, J. D. 2016. Downregulation of let-7b promotes COL1A1 and COL1A2 expression in dermis and skin fibroblasts during heat wound repair. Molecular Medicine Reports, 13, 2683–2688.

Maffulli, N., Moller, H. D. & Evans, C. H. 2002. Tendon healing: can it be optimised? Br J Sports Med, 36, 315–6.

Mazzone, M. F. & Mccue, T. 2002. Common conditions of the achilles tendon. Am Fam Physician, 65, 1805–10.

Myhrvold, S. B., Brouwer, E. F., Andresen, T. K. M., Rydevik, K., Amundsen, M., Grun, W., Butt, F., Valberg, M., Ulstein, S. & Hoelsbrekken, S. E. 2022. Nonoperative or Surgical Treatment of Acute Achilles’ Tendon Rupture. N Engl J Med, 386, 1409–1420.

Oh, S. J., Erb, H. H., Hobisch, A., Santer, F. R. & Culig, Z. 2012. Sorafenib decreases proliferation and induces apoptosis of prostate cancer cells by inhibition of the androgen receptor and Akt signaling pathways. Endocr Relat Cancer, 19, 305–19.

Ong, C. T., Khoo, Y. T., Mukhopadhyay, A., Masilamani, J., Do, D. V., Lim, I. J. & Phan, T. T. 2010. Comparative proteomic analysis between normal skin and keloid scar. British Journal of Dermatology, 162, 1302–1315.

Pires Da Silva, J., Monceaux, K., Guilbert, A., Gressette, M., Piquereau, J., Novotova, M., Ventura-Clapier, R., Garnier, A. & Lemaire, C. 2020. SIRT1 Protects the Heart from ER Stress-Induced Injury by Promoting eEF2K/eEF2-Dependent Autophagy. Cells, 9.

Qu, F., Pintauro, M. P., Haughan, J. E., Henning, E. A., Esterhai, J. L., Schaer, T. P., Mauck, R. L. & Fisher, M. B. 2015. Repair of dense connective tissues via biomaterial-mediated matrix reprogramming of the wound interface. Biomaterials, 39, 85–94.

Ren, H., Zhao, F., Zhang, Q., Huang, X. & Wang, Z. 2022. Autophagy and skin wound healing. Burns Trauma, 10, tkac003.

Rose, A. J., Alsted, T. J., Jensen, T. E., Kobbero, J. B., Maarbjerg, S. J., Jensen, J. & Richter, E. A. 2009. A Ca(2+)-calmodulin-eEF2K-eEF2 signalling cascade, but not AMPK, contributes to the suppression of skeletal muscle protein synthesis during contractions. J Physiol, 587, 1547–63.

Sagnol, S., Yang, Y., Bessin, Y., Allemand, F., Hapkova, I., Notarnicola, C., Guichou, J. F., Faure, S., Labesse, G. & De Santa Barbara, P. 2014. Homodimerization of RBPMS2 through a new RRM-interaction motif is necessary to control smooth muscle plasticity. Nucleic Acids Res, 42, 10173–84.

Saraswati, S., Marrow, S. M. W., Watch, L. A. & Young, P. P. 2019. Identification of a pro-angiogenic functional role for FSP1-positive fibroblast subtype in wound healing. Nat Commun, 10, 3027.

Saxena, S., Vekaria, H., Sullivan, P. G. & Seifert, A. W. 2019. Connective tissue fibroblasts from highly regenerative mammals are refractory to ROS-induced cellular senescence. Nat Commun, 10, 4400.

Schwanhausser, B., Busse, D., Li, N., Dittmar, G., Schuchhardt, J., Wolf, J., Chen, W. & Selbach, M. 2011. Global quantification of mammalian gene expression control. Nature, 473, 337–42.

Shah, J. M., Omar, E., Pai, D. R. & Sood, S. 2012. Cellular events and biomarkers of wound healing. Indian J Plast Surg, 45, 220–8.

Shi, N., Chen, X., Liu, R., Wang, D., Su, M., Wang, Q., He, A. & Gu, H. 2018. Eukaryotic elongation factors 2 promotes tumor cell proliferation and correlates with poor prognosis in ovarian cancer. Tissue Cell, 53, 53–60.

Svedman, S., Juthberg, R., Edman, G. & Ackermann, P. W. 2018. Reduced Time to Surgery Improves Patient-Reported Outcome After Achilles Tendon Rupture. Am J Sports Med, 46, 2929–2934.

Tchoukalova, Y. D., Zacharias, S. R. C., Mitchell, N., Madsen, C., Myers, C. E., Gadalla, D., Skinner, J., Kopaczka, K., Gramignoli, R. & Lott, D. G. 2022. Human amniotic epithelial cell transplantation improves scar remodeling in a rabbit model of acute vocal fold injury: a pilot study. Stem Cell Res Ther, 13, 31.

Tracy, L. E., Minasian, R. A. & Caterson, E. J. 2016. Extracellular Matrix and Dermal Fibroblast Function in the Healing Wound. Adv Wound Care (New Rochelle), 5, 119–136.

Tsuchiya, K. 2021. Switching from Apoptosis to Pyroptosis: Gasdermin-Elicited Inflammation and Antitumor Immunity. Int J Mol Sci, 22.

Vescarelli, E., Pilloni, A., Dominici, F., Pontecorvi, P., Angeloni, A., Polimeni, A., Ceccarelli, S. & Marchese, C. 2017. Autophagy activation is required for myofibroblast differentiation during healing of oral mucosa. Journal of Clinical Periodontology, 44, 1039–1050.

Wilhelm, M., Schlegl, J., Hahne, H., Gholami, A. M., Lieberenz, M., Savitski, M. M., Ziegler, E., Butzmann, L., Gessulat, S., Marx, H., Mathieson, T., Lemeer, S., Schnatbaum, K., Reimer, U., Wenschuh, H., Mollenhauer, M., Slotta-Huspenina, J., Boese, J. H., Bantscheff, M., Gerstmair, A., Faerber, F. & Kuster, B. 2014. Mass-spectrometry-based draft of the human proteome. Nature, 509, 582–7.

Wong, T., Mcgrath, J. A. & Navsaria, H. 2007. The role of fibroblasts in tissue engineering and regeneration. Br J Dermatol, 156, 1149–55.

Xie, Z. Y., Xiao, Z. H. & Wang, F. F. 2018. Inhibition of autophagy reverses alcohol-induced hepatic stellate cells activation through activation of Nrf2-Keap1-ARE signaling pathway. Biochimie, 147, 55–62.

Yang, Y., Lin, Z., Cheng, J., Ding, S., Mao, W. W., Shi, S., Liang, B. & Jiang, L. 2020. The roles of autophagy in osteogenic differentiation in rat ligamentum fibroblasts: Evidence and possible implications. FASEB J, 34, 8876–8886.

Yao, Q., Liu, B. Q., Li, H., Mcgarrigle, D., Xing, B. W., Zhou, M. T., Wang, Z., Zhang, J. J., Huang, X. Y. & Guo, L. 2014. C-terminal Src kinase (Csk)-mediated phosphorylation of eukaryotic elongation factor 2 (eEF2) promotes proteolytic cleavage and nuclear translocation of eEF2. J Biol Chem, 289, 12666–78.

Zhang, L., Wang, K., Lei, Y., Li, Q., Nice, E. C. & Huang, C. 2015. Redox signaling: Potential arbitrator of autophagy and apoptosis in therapeutic response. Free Radic Biol Med, 89, 452–65.

